# Maternal vitamin D deficiency impairs heart formation in mouse offspring through a change in 3D-chromatin structure

**DOI:** 10.1101/2020.12.17.423263

**Authors:** Eva M. Seipelt, Paul Bensadoun, Satish Sati, Charlène Couturier, Julien Astier, Lourdes Mounien, Jean-François Landrier, Michel Pucéat

## Abstract

The origins of congenital heart diseases, the most common congenital diseases are still largely unknown. Environmental factors are now emerging as major causes of these diseases. Vitamin D deficiency has become a public health burden, notably for childbearing age, pregnant and breastfeeding women. Since maternal 25-hydroxyvitamin D (25(OH)D) determined fetal and neonatal 25(OH)D status, foetuses exposed to insufficient levels of vitamin D, may feature developmental defects.

Herein, we investigated the effects of maternal vitamin D deficiency on cardiovascular defects in early and later life of offsprings in two generations as well as the molecular mechanisms underlying vitamin D effect.

Eight weeks before and during pregnancy, C57BL/6JRj female mice received a sufficient or vitamin D deficient diet ((1.0 IU/g in control vs 0.0 IU/g in Vitamin D Deficient (VDD) group). E16.5 Embryos of maternal VDD diet featured hypertrophic heart revealed by a thicker left ventricular (LV) wall and septum. RNAseq analysis of LV revealed 1555 transcripts differentially expressed in the VDD group and among them cardiac transcription factors and constitutive cardiac genes (*tbx5, gata4, myl2*). Anti-Vitamin D receptor (VDR) Chip-seq from chromatin of E16.5 LV uncovered different targeting of *tbx5* and *tbx3* loci by VDR in the VDD vs control embryos. Anti-CTCF ChIP-loop experiments focusing on the *Tbx3* and *Tbx5* loci uncovered a change in the Topology Associated Domains associated with these loci.

Echocardiography of 2-months-old VDD offspring revealed a significantly thicker left ventricle and increased fractional shortening while 6-months-old mice featured cardiac decompensation and in turn failing LV.

Maternal vitamin D deficiency severely affects heart formation following a change in chromatin conformation on cardiac gene loci and impacts function of adult hearts in two generations. These defects are likely to be at the origin of cardiovascular diseases in the adulthood.

## INTRODUCTION

Congenital heart diseases (CHD) are the most common congenital malformations at birth (Hoffman and Kaplan, 2002) and result from defect in the cardiogenic transcriptional program (Zaidi and Brueckner, 2017) as well as from epigenetic dysregulation of the later (Moore-Morris et al., 2018). The prevalence of CHD is 8 per 1000 live birth(Hoffman and Kaplan, 2002). As techniques of corrective and palliative surgery continuously improve for last three decades, the number of new-born becoming adult has considerably increased.

The likely multifactorial causes of CHD, are still poorly known (Gelb, 2015). The growing population of adult CHD patients calls for a better understanding of the origins of these diseases.

Maternal nutrition impacts normal foetal development and may predispose the offspring to future cardiovascular diseases (CVD) (Hanson and Gluckman, 2014). During the past decades, maternal vitamin D insufficiency has become a worldwide issue since approximately 54% of pregnant women present an impaired vitamin D status with a 25-hydroxyvitamin D concentration below 50 nmol/L (Saraf et al., 2016).

Furthermore deficit in vitamin D is interrelated with obesity (Paschou et al., 2019) which represents two pandemic conditions of high risk for CHD.

Vitamin D is a fat soluble steroid and nuclear hormone either ingested through the diet or produced *de novo* from 7-dehydrocholesterol in the skin after exposure to ultraviolet B radiation from sunlight (Christakos et al., 2016).

Vitamin D or its metabolites are involved in adult CVD. Animal and *in vitro* studies showed that in absence of VDR (Knockout models) or in animal fed with vitamin D depleted diet, the function of adult heart is impacted (Assalin et al., 2013; Rahman et al., 2007; Reddy Vanga et al., 2010; Xiang et al., 2005). Data from the Offspring Cohort of the Framingham Heart Study have revealed a correlation between vitamin D levels and CVD (Wang et al., 2008).

While vitamin D plays an important role in adult heart homeostasis, whether maternal vitamin D may effect heart formation of the offspring has not been investigated.

Recently in a population-based case-control family study, it has been reported that a compromised maternal vitamin D status is associated with an approximately two-fold increased prevalence of congenital heart defects in offspring (Koster et al., 2018). Another study reported that the maternal vitamin D status seems to be inversely correlated to the risk of CHD in the child (Mokhtar et al., 2019). Nevertheless, if direct, how deficiency in maternal vitamin D impacts embryonic heart formation is still not documented and the molecular mechanisms underlying the effect of vitamin D on heart formation are not known.

The aims of the current study were to evaluate the impact of maternal vitamin D on mouse cardiogenesis and to uncover the molecular mechanisms mediating the effects of vitamin D deficiency.

We found that absence of vitamin D during pregnancy leads to hypertrophy of the left ventricle and septum of embryos. The hypertrophy was still observed in adults. The process was accompanied by a dysregulation of the embryonic cardiac genetic programme following a change in 3D chromatin structure.

## RESULTS

### Maternal vitamin D deficiency leads to left ventricular hypertrophy in the embryo

We first monitored the status of maternal vitamin D in female fed with a vitamin D-null diet. Mouse body weight and food intake were not impacted by the Vitamin D deficient (VDD) diet (Figure S1 AB). 25-hydroxivitamin D concentrations were below the detection limit of the assay for the VDD mother compared to controls (Figure S1C).Maternal VDD diet consumption prior and during gestation led to significant smaller weaning (4 weeks old) body weight in mice offspring (Figure S1D).

Maternal VDD diet consumption prior to and during gestation led to significant thicker left ventricle (LV) wall and septum in the embryonic E16.5 hearts compared to those from control mice (Figure 1 A-D). There was no statistical difference between the two conditions for the right ventricle wall (Figure 1E). The size of cardiomyocytes was visualized using FITC conjugated WGA and imaged in confocal microscopy. Cardiomyocytes from VDD left ventricle hearts were significantly larger (Figure 1 G) when compared to the controls (Figure 1 F-H).

**Fig 1.**
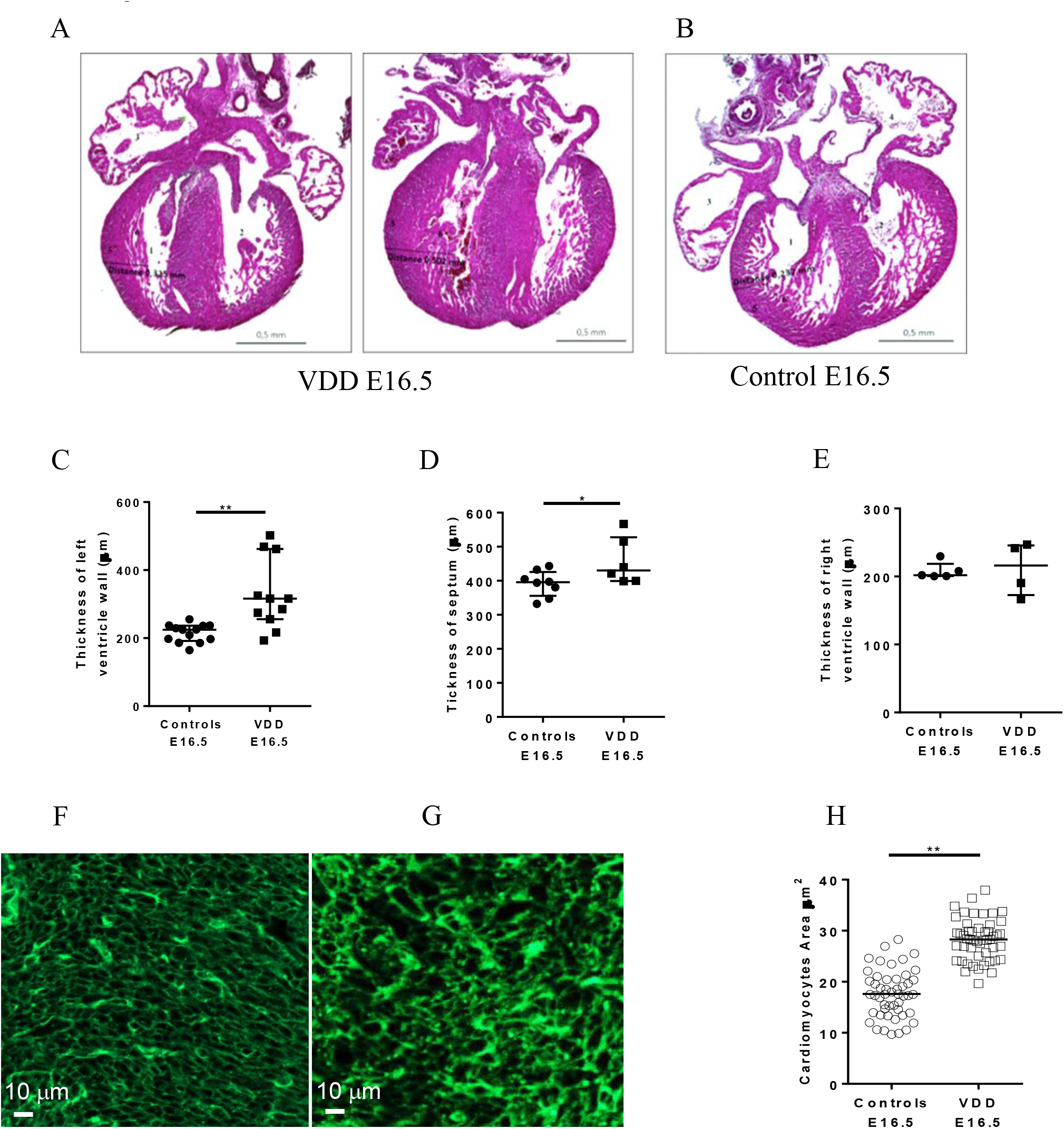
Cardiac morphology of E16.5 mice offspring from vitamin D deficient (VDD) or control mothers. Eosin-hematoxylin slide of (A) VDD hearts and (B) Control heart.(C) Graphs of the left ventricle wall, (D) septum and (E) right ventricular wall of control and VDD embryos (** p< 0.01, * p< 0.05). (F) Cardiomyocytes area from control and (G) VDD embryo. Left ventricular sections were stained with FITC-WGA and measured using NIH image. (H) graph illustrating measurements in 50 to 100 cells (** P < 0.01)

### Gene Transcripts from vitamin D deficient embryos were dysregulated during cardiogenesis

To get insight on the mechanisms underlying ventricular hypertrophy, we interrogated the cardiac (LV) transcriptome of embryonic E16.5 hearts. RNA-seq analysis using 8 left ventricles revealed 1555 transcripts differentially expressed in the VDD group with 1058 genes up regulated (FC > 1.5) and 497 genes downregulated (FC < −1.5) (heatmap Figure 2A, supplemental data 1)

**Fig 2.**
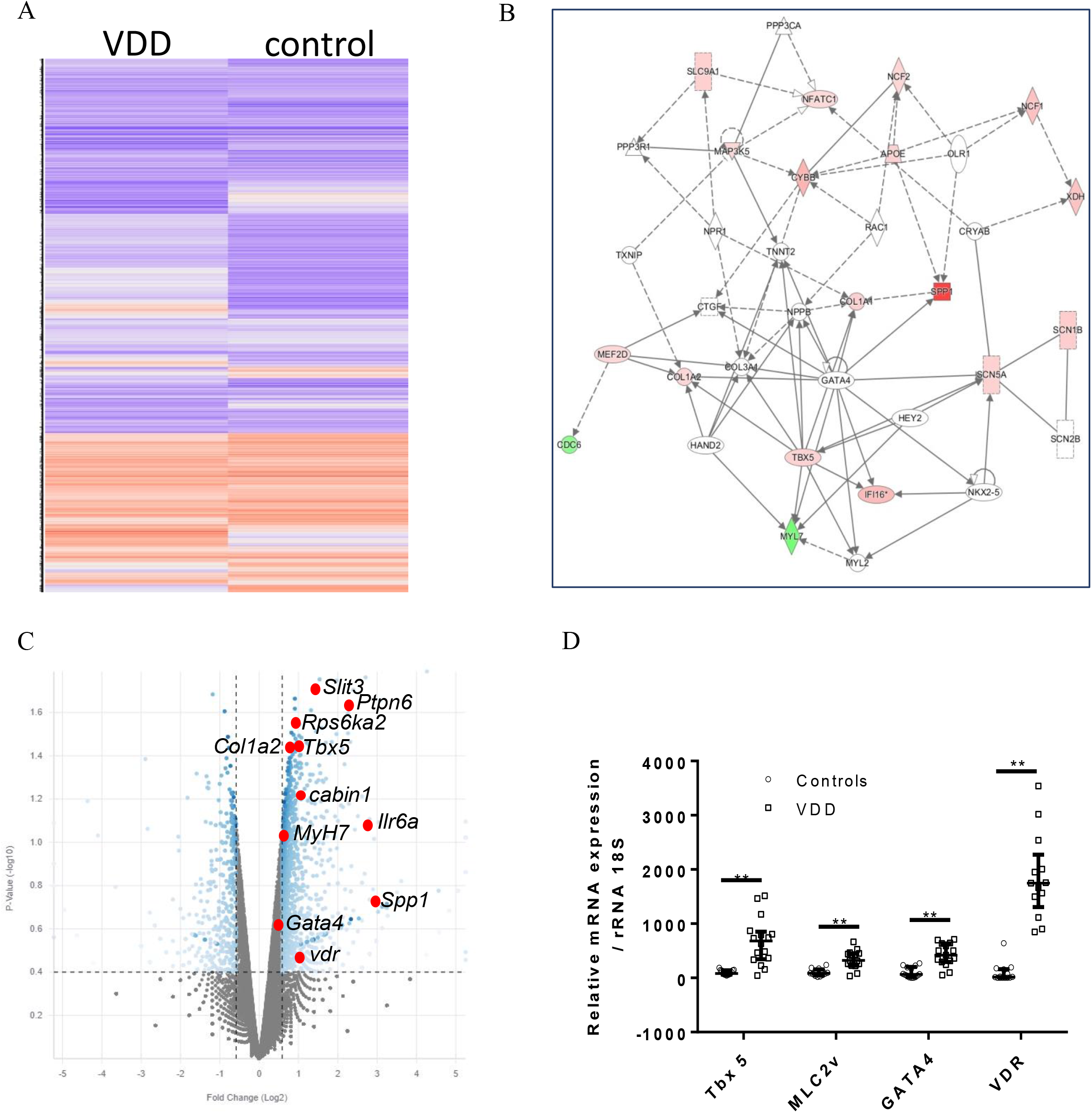
Transcriptomic analysis of E16.5 hearts. (A) heatmap of genes expressed in VDD or control E16.5 left ventricle and septum; blue indicates lower and orange higher expression (B) Expression of RNA transcripts implicated in the cardiogenesis were dysregulated in the E16.5 heart either under-regulated (green) or over-regulated (red) using the data from RNA-sequencing on Ingenuity Pathway Analysis software.(C) volcano plot highlighting upregulated genes in VDD versus control left entricle and septum of E16.5 embryos (D) Real time Q-PCR of *Tbx5, Gata4, Mlc2v* and *Vdr* (** p< 0.01)

A pathway analysis revealed that genes upregulated in VDD were mostly involved in embryonic development and more specifically immune system and cardiac development, response to cytokine stimulus and chromatin modification (supplemental data 2). Figure 2B highlights IPA generated network with genes upregulated (pink boxes) in VDD hearts regulating formation of the heart or size of the ventricle.

A Volcano plot illustrated in figure 2C points another set of genes among those highly up-regulated and involved in heart hypertrophy such as *Slit3, Ptpn6, Rps6ka2, Myh7 and Il6Ra as well as cabin 1* (Figure 2C).

Upregulation of a few cardiac specific genes *tbx5, gata4 and myl2* in VDD embryonic hearts were validated by real-time PCR (Figure 2 D). *Tbx5, gata4, myl2* were all overexpressed (P <0.05) in VDD hearts compared to control. The *VDR* was also upregulated in VDD hearts

### Gene regulatory regions were differentially targeted by VDR when exposed to maternal vitamin D deficiency

To identify potential loci differentially targeted by the Vitamin D Receptor (VDR) we performed Immunoprecipitation of chromatin of E16.5 left ventricle using an anti-VDR antibody. A genome wide view of Chip-seq analysis revealed that many gene regulatory regions occupied by VDR in control heart were lost in VDD hearts (supplemental Figure 2; ChIP-peaks in supplemental data 3).

*Tbx5* is required for left ventricle as well as septum formation (Takeuchi et al., 2003), the two regions affected in VDD embryonic hearts. We thus next focused on a specific genomic region including *Tbx3* and *Tbx5* (Figure 3). *Tbx5* and *Tbx3* loci were differentially occupied by VDR in the VDD embryonic hearts when compared to control hearts (Figure 3A). Enrichment peaks were present in control ventricles all over the locus forming topology associated domains (van Weerd et al., 2014) (Figure 3B). Interestingly, four main peaks (arrows in Figure 3A) were CTCF-enriched regions (van Weerd et al., 2014) and were lost in VDD mouse left ventricles.

**Fig 3.**
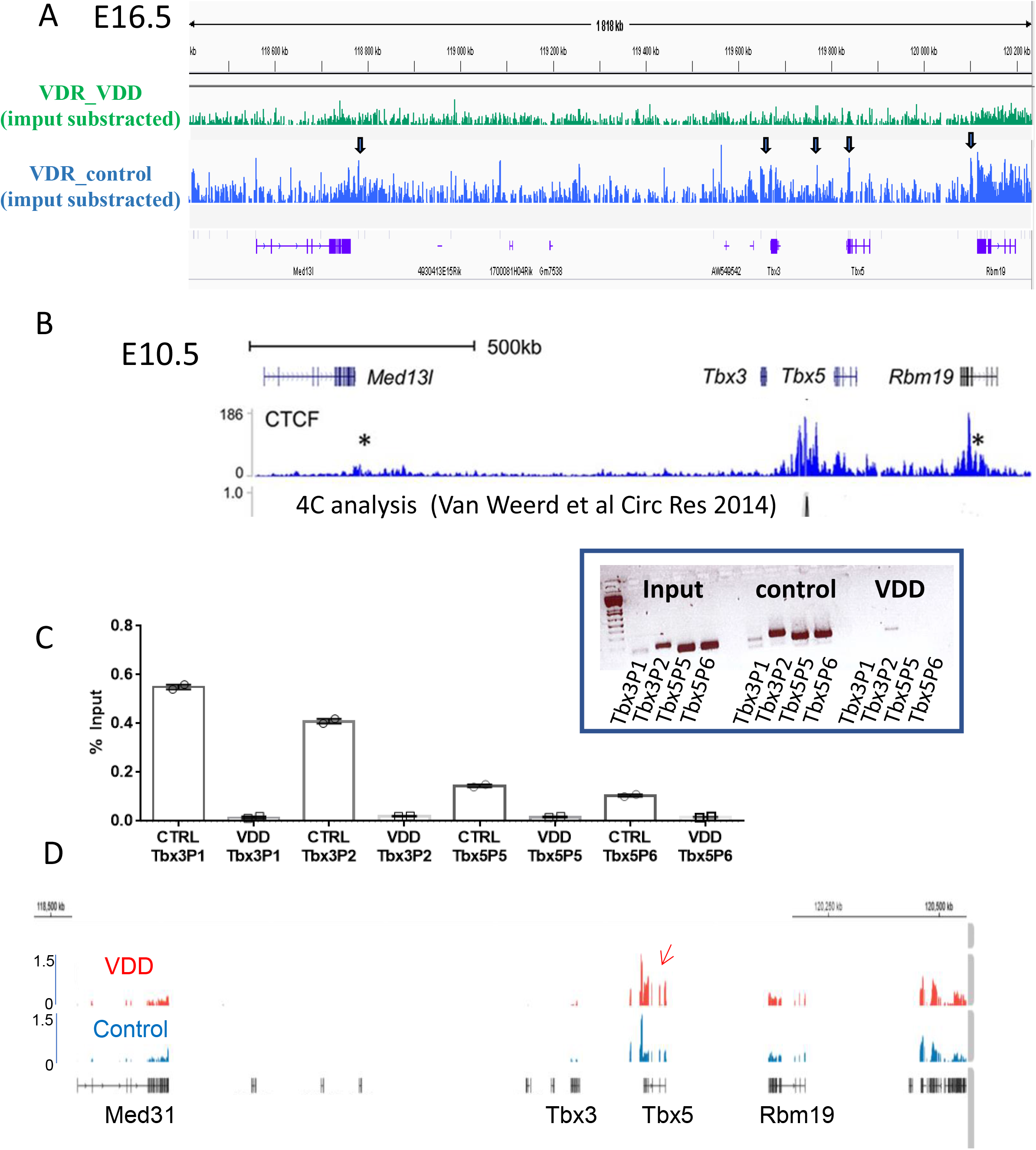
Chip sequencing analysis of VDR-targeted genes in the E16.5 hearts (left ventricle and septum) within the *Tbx3/Tbx5* locus (A) Enrichment peaks over input in control or VDD hearts. (B) 4C analysis of the locus (data from Van Weerd et al^19^) (C) ChIP QPCR validation of *Tbx3* and *Tbx5* promoter occupancy by VDR. The experiment was performed in duplicate. Inset gel of the PCR amplicons. (D) Highlight of the RNA-sequencing peaks within the *Tbx3/Tbx5* locus.

We then used ChIP-QPCR to validate the binding domains of VDR within the *Tbx3 /Tbx5* loci. Figure 3C illustrates the enrichment over input of VDR on *Tbx3* and *Tbx5* promoter regions. The enrichment on all 4 genomic regions in control E16.5 embryonic LV was fully lost in hearts of embryos of VDD mothers. This was confirmed by running the amplicons on gels (Figure 3C inset). The loss in VDR enrichment on *Tbx5* locus correlated with a de-repression of the locus and in turn, an increase in *Tbx5* expression as shown by RNA-sequencing focused on this specific locus (Figure 3D).

Encode Data of occupancy of different histone marks on this locus in E16.5 whole hearts revealed the presence of both activating marks usually present on enhancers (H3K4me1/2) or promoter (H3K4ac2) as well as repressive marks (H3K27me3). Occupancy by H3K36me3 indicating RNA pol2 activity was low on Tbx5 locus (Supplemental Fig 3). Altogether, this epigenetic landscape reflects a pause status of the *Tbx5* promoter at this stage of heart development.

#### Anti-CTCF ChIP and Chromosome Configuration Capture of tbx3/tbx5 locus

To get further insight into the epigenetic landscape of the *tbx3/tbx5* loci, we designed a ChIP-loop experiment. Chromatin was prepared from LV of E16.5 control heart or VDD hearts. CTCF was immune-precipitated and the chromatin structure surrounding DNA bound to CTCF was monitored by a 3-C approach.

Two main CTCF sites making a boundary between the *tbx3* and *tbx5* loci were previously identified (van Weerd et al., 2014). We thus interrogated these regions using a ChIP-loop assay.

C-loop revealed that the DNA contacts at the CTCF boundary regions between *tbx3* and *tbx5* locus were absent in control E16.5 hearts but gained in VDD hearts (Figure 4A and inset). Indeed PCR primers flanking a region of 29 kb making a loop bridging at the CTCF1 site between a region upstream of *tbx5* and a region downstream of *med31*, could amplify a short sequence in VDD hearts indicating formation of a TAD that links together the DNAs (Figure 4A inset). Interestingly, PCR primers flanking a region of 432 kb also amplified a short DNA region as an index of DNA interaction at the CTCF site 2 bridging the two loops between a region upstream of *rbm19* and a region downstream of *med31* (Figure 4A, inset).

**Fig 4.**
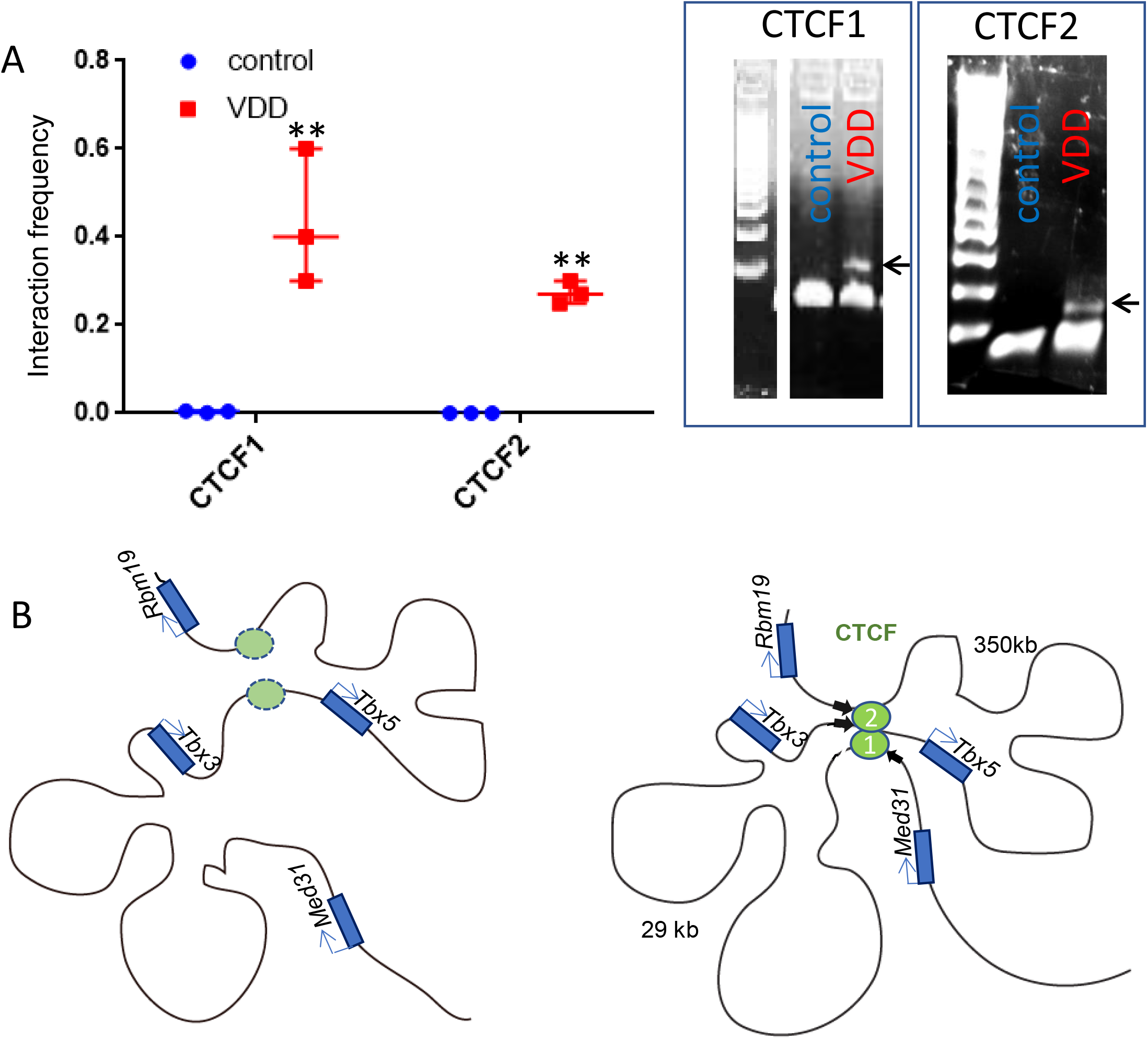
Anti-CTCF ChIP-loop from chromatin extracted from left ventricle and septum of E16.5 embryonic hearts. (A) Relative abundance of CTCF in CTCF-immunoprecipitated DNA. Experiment was repeated 3 times and PCR repeated 3 times (** p< 0.01); inset shows the amplicon on gel. (B) cartoon showing the 3D configuration landscape of the *Tbx3/Tbx5* locus (inspired from Van Weerd et al 2014) in control (left panel) and VDD condition (right panel). Black arrows indicate positions of 3C-PCR primers

Figure 4B shows the topology associated domains within the *tbx3/tbx5* loci as previously described in E10.5 embryonic hearts (van Weerd et al., 2014). The CTCF1-CTCF2 bridging TADs were not formed in E16.5 hearts but reformed in heart from embryos of vitamin D deficient mothers.

#### Maternal vitamin D deficiency programs cardiac dysfunction in adult offspring

To explore the long-term effect of maternal vitamin D deficiency in the offspring we used echocardiography to evaluate the cardiac function at 2-months and 6-months of age. 2-months-old VDD males still featured a significantly thicker left ventricle wall and increased fractional shortening compared to the control (*P* < 0.01) (Fig 5B and 5C).

**Fig 5.**
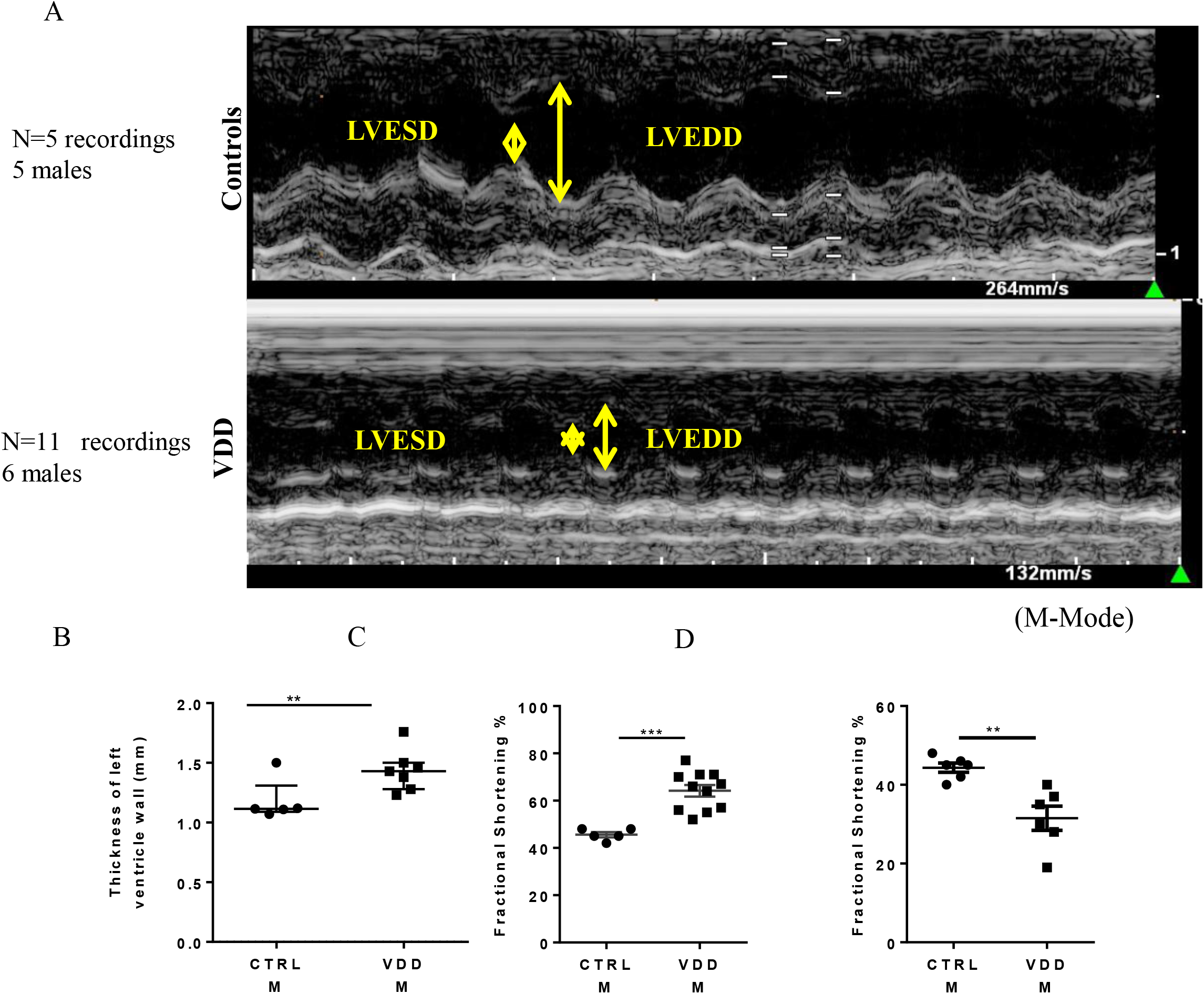
Cardiac morphology and function of 2-months and 6-months-old male offspring. (A) Echocardiography of 2-months-old offspring. (B) Thickness of left ventricle wall (** *p*< 0.01), (C) fractional shortening of 2 months old control and VDD hearts (**p< 0.01) (D) Fractional shortening of 6 months old control and VDD hearts (** *p*< 0.01). LVESD: left ventricular end systolic dimension, LVEDD: left ventricular end diastolic dimension. The echocardiography was performed in 5 to 10 mice

6-months-old VDD males presented a decreased fractional shortening (P < 0.01) (Fig 5D).

#### Maternal vitamin D deficiency-induced cardiac dysfunction in adult offspring is observed in the second generation

As we identified an epigenetic cause of embryonic cardiac hypertrophy that was maintained in the adult, we investigated cardiac function in the second generation. We bred adult females originally born from VDD mothers with males. The offspring of this breeding set up were left to grow up to two months adult age. Echocardiography revealed an increase in shortening fraction and an increase in left ventricular wall thickness indexes of ventricular hypertrophy (figure 6)

**Figure 6:**
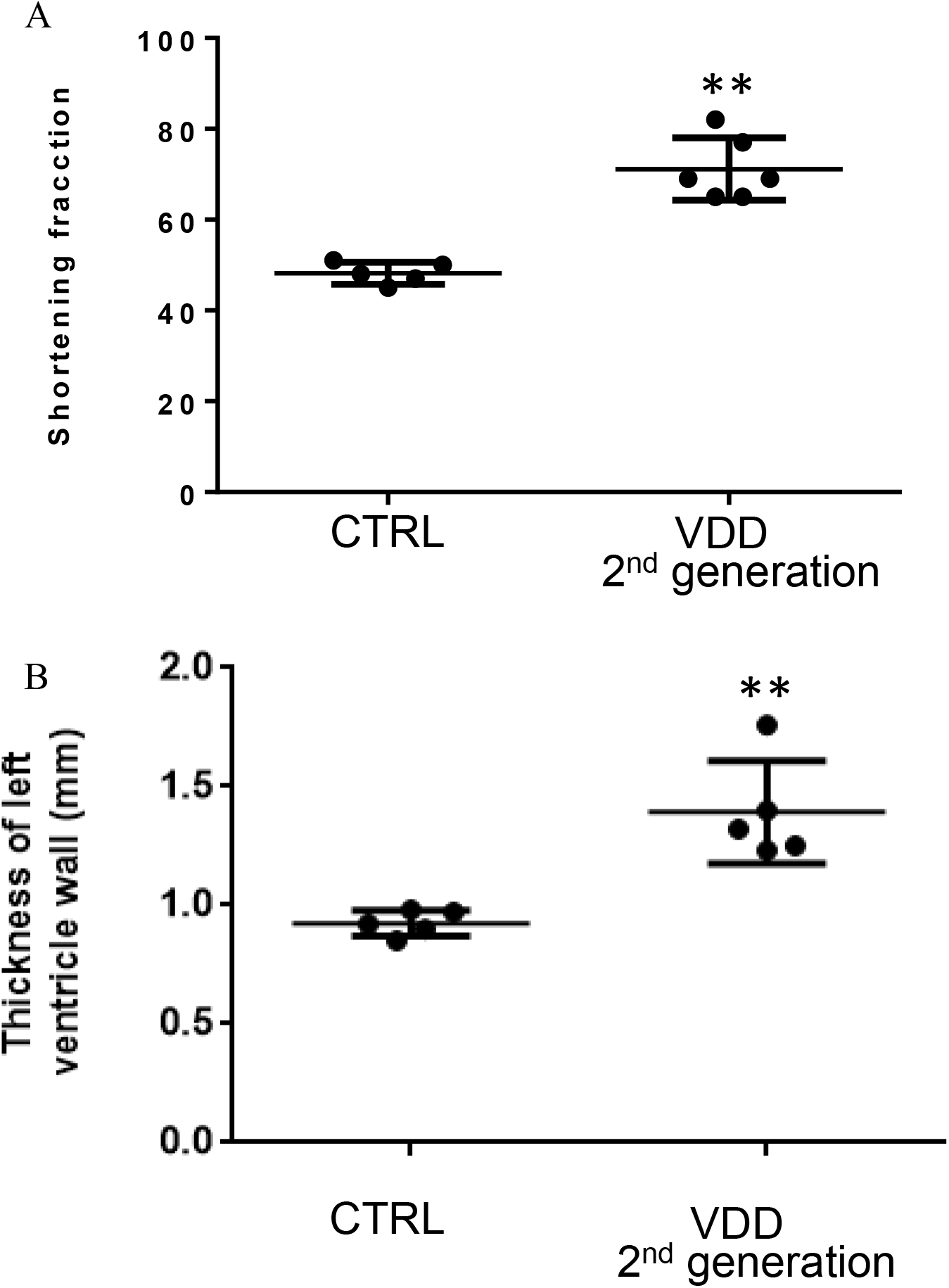
Echocardiography of second generation mice. A, left ventricle shortening fraction. B: left ventricular wall thickness

## Discussion

Among environmental factors, maternal nutrition may play a major role in proper embryonic cardiac development and in turn in CHD and may affect cardiovascular health of the offspring at adulthood.

Herein, we examined the effects of maternal vitamin D deficiency (VDD), a nuclear hormone exerting pleiotropic effects (Christakos et al., 2016) on the potential incidence of CHD in the developing embryo and long-term consequences on cardiac function in the offspring. We found that embryonic E16.5 mouse heart featured both left ventricular and septum hypertrophy after a maternal VDD induced by diet during the antenatal period (8 weeks before mating until birth). The left ventricular hypertrophy was still observed at 2 months-of age in the offspring even if the offspring was not exposed to VDD after birth. Furthermore, the second generation of mice fed like their parents with a regular vitamin D containing diet still featured cardiac hypertrophy at the adulthood. Altogether, these observations suggest that VDD triggered transgenerational epigenetic changes possibly through somatic cells including maternal mitochondria and/or germ cells.

The early compensatory hypertrophy expected from adaptation of the embryo to a stressing nutritional environment leads to maladaptive hypertrophy and in turn cardiac failure as observed in the first generation 6-months-old VDD males. These data point to a strict requirement of maternal vitamin D for healthy cardiac functions of the offspring. In contrast to the zebrafish, embryonic hearts from VDD did not feature any change in cell cycle genes such as erbb2, *ccnd1*, e2f, or *mitf* found upregulated in VD overexpressing fish hearts (Han et al., 2019).

Our data thus significantly extend the observation of Gezmish and collaborators who reported a cardiac hypertrophy in 21 days adult rat offspring from mother with depleted diet in vitamin D compared to those from supplemented vitamin D diet (Gezmish et al., 2010). Furthermore, our observations open the path toward an epigenetic printed mechanism taking place during embryonic development and underlying the long term effects of VDD.

Transcriptomic analysis of left ventricular hearts, revealed a dysregulation of major biological pathways in embryonic hearts. These included embryonic developmental pathways. More specifically, major transcription factors playing at early stages of cardiogenesis key roles in determination of cardiac lineages such as *Tbx5* or *Gata4* were up-regulated at embryonic stage E16.5 as well as some genes involved in the process of cardiac hypertrophy such as *Rps6ka2* (Li et al., 2013), *Slt3* (Gong et al., 2020) and *Ptnp6* involved in angiotensin AT1/EGF pro-hypertrophic signalling pathway (Kagiyama et al., 2002) and a direct target of VDR (Kumar et al., 2020). Calbin1 which inhibits the pro-hypertrophic calcineurin pathway (De Windt et al., 2001) was also upregulated likely to counteract the hypertrophic response of VDD hearts. Thus the hypertrophic phenotype was primed during formation of the heart.

VDR is often co-localised with the cohesin complex (Lu et al., 2018) and in 15% of sites specifically with cohesin/CTCF complex (Parelho et al., 2008). We first interrogated the whole genome of embryonic E16.5 left ventricle from mother fed with a regular or a VDD diet, as to the VDR binding sites. Immunoprecipitation of the VDR revealed that the VDR-enrichment of many loci were lost in VDD hearts.

More specifically we looked at the *Tbx5* locus. VDR occupancy was different in both *Tbx5* and *Tbx3* regulatory regions in the VDD embryos when compared to the controls. The enrichment of VDR on CTCF sites within these gene loci as identified by Van Weerd et al (van Weerd et al., 2014) was specifically lost in VDD hearts.

The explanation for this observation could be several. The nuclear translocation, deposit or at least the high affinity binding of VDR on genomic loci depends upon the binding of vitamin D in line with the requirement of the ligand to ensure the transcriptional activity (Christakos et al., 2016; Nagpal et al., 2005). Alternatively, Vitamin D may regulate CTCF binding to DNA in a non VDR-dependent manner (Seuter et al., 2016). This could explain why we observed cardiac hypertrophy in our embryos deficient in vitamin D but featuring an upregulation of VDR while Chen and collaborators reported that deletion of VDR mRNA was also associated with an induction of myocyte hypertrophy either *in vitro* or *in vivo* in adult (Chen et al., 2019). Interestingly even if overexpressed as in our VDD embryonic hearts VDR in the absence of vitamin D could not translocate into the nucleus thus preventing any nuclear function which is also in line with the hypertrophic phenotype of VDR KO adult hearts. Whether other ligands of VDR (bile acids, melatonin) (Fang et al., 2020; Han et al., 2010) could compensate for the absence of vitamin D is unlikely in cardiomyocytes.

We thus hypothesised that gene upregulation could be linked to a change in 3D-chromatin structure and more specifically CTCF mediated DNA loops. We choose the *Tbx3/Tbx5* loci taking into account the cardiac phenotype of embryos and as enriched in CTCF sites including two main ones that isolate the two genes (van Weerd et al., 2014) into separate DNA loops, a condition likely required at E10.5 stage of development featuring high *Tbx5* expression (van Weerd et al., 2014) *.Tbx5* starts to be downregulated at E16.5 under normal conditions while it was upregulated in VDD hearts. Anti-CTCF ChIP-3C revealed that the two DNA strands belonging to the separate loops interacted in VDD hearts but not in control hearts. This indicates that under control diet condition, the VDR/vitamin D complex associated with CTCF sites likely disrupts the cohesin complex and unfolds the loops forming the TADs. This in turns impairs Tbx5 expression as a gene near a boundary thus highly sensitive to cohesin loss (Luppino et al., 2020). In contrast, in the absence of the complex, the interaction is regained and *Tbx5* which should have remained silenced is re-expressed.

We thus report a direct effect of Vitamin D and its receptor on the chromatin 3D structure. We focused on the *Tbx5* locus, a key gene to specify both left ventricle and septum (Takeuchi et al., 2003), which acts in a dosage dependent manner. Tbx5 haplo-insufficiency such as in Holt-Oram syndrome is responsible for cardiac defects while overexpression specifically of the long isoform expressed from E13.5 in embryonic heart leads to cardiac hypertrophy (Georges et al., 2008). Although the VDR/vitamin D complex features a broad range of occupancy sites within the full genome, the embryos featured normal growth and besides hearts, no macroscopic defects was observed for any other organs. However, we cannot exclude that VDD by triggering changes in chromatin structure may increase cell plasticity and affect specification or determination of other cell lineages.

For example as we recently reported, maternal VDD likely affects the identity and/or function of the adipocytes and primes future metabolic disorders in the offsprings when reaching adulthood (Seipelt et al., 2020). Embryonic cardiac metabolism may also be affected in utero specifically when cardiomyocytes switch from glycolytic to oxidative metabolism at E11.5-E13.5 and could further contribute to the cardiac hypertrophic phenotype.

Maternal vitamin D thus turns out to be a key nutriment whose concentration should be tightly controlled to ensure a proper heart formation and to prevent CHD and future acquired cardiovascular diseases. Its requirement together VDR to maintain a proper chromatin structure makes vitamin D even more important as a regulator of metabolism to prevent any cardiac or metabolic disease with transgenerational transmission (Sales et al., 2017).

## Material and Methods

### Animal Experiments

The protocol received the agreement of Aix-Marseille University Ethics Committee and the French Ministry of Research (APAFIS#1300-2015072112279135). Eight-week-old female and male C57BL/6JRJ mice were obtained from Janvier Labs (Le Genest-Saint-Isle, France), fed ad libitum with control food (chow diet A04 from Safe-diets, Augy France) during the 1-week acclimation period and with full access to drinking water. The animals were maintained at 22°C under a 12-hour light, 12-hour dark cycle and a 20% humidity level. Female mice (15 per group) were assigned into one of the two experimental groups depending on the diet i.e. control (AIN-93G with vitamin D3, 1.0 IU/g) or vitamin D-depleted (AIN-93G without vitamin D3, 0.0 IU/g) for eight-weeks. Females were mated with males. Weight gain was measured once a week and dietary at 3-weeks of pre-mate diet, at 5 days and 15 days of gestational stage (Supplemental data Figure 1). Females were euthanized by cervical dislocation and hearts of E16.5 embryos were collected. Left Ventricle (LV) was dissected out for RNA sequencing, anti-VDR ChIP-sequencing and ChIP-3C. Whole hearts were fixed with paraformaldehyde 4% overnight and later used for histology. After delivery, all remaining females and their offspring were fed with control diet (AIN-93G) until the study (6months-old offspring). Only males were used for the experiments.

### Biochemical analysis

Maternal vitamin D status 25(OH)D serum concentrations were measured using an in vitro diagnostic enzyme immunoassay kit 25-OH Vitamin D (direct) ELISA kit (PromoKine) according to the manufacturer’s protocol.

### Heart Histology

E16.5 hearts (n = 11 for VDD and n = 13 for controls) were fixed with 4% paraformaldehyde over night at 4°C, dehydrated with 50%, 90% and 100% ethanol over 4 hours. Then hearts were embedded with paraffin. 8μm sections of the heart were stained with Harris Hematoxylin Solution Modified (Sigma Aldrich) and Eosin y alcoholic solution (Sigma Aldrich). Then, the slides were mounted with Eukitt Quick hardening (Sigma Aldrich). Heart wall thickness was determined using the average of values obtained from three consecutive sections for each heart using ZEN software (Zeiss).

### Immunofluorescence

Hearts were embedded with OCT-sucrose (25%sucrose). Each sections of heart were 8μm. Slides were washed with PBS 1X, then placed in PBS-Triton 0.1% for 10min. Slides were blocked with 4.5mL PBS-Triton X100 0.1%, 0.1g BSA with WGA, Alexa Fluor 488 conjugate (Invitrogen) and DAPI 1X solution following the manufacturer’s protocols. Slides were washed with PBS 1X and mounted with fluoromount mounting medium (Southern-Biotec). For each heart, we scored the area of at least 100 cardiomyocytes in 3 different regions of the ventricle from apex to atrium. Cell size was calculated after thresholding the image using NIH image.

### RNA extraction real time PCR and RNA sequencing

Total RNA was extracted from E16.5 LV hearts (VDD n=8 and Controls n=8) using Zymo Research Corp kit ZR RNA Miniprep following the manufacturer’s protocol. For the real time PCR, one μg of total RNA from LV was used to synthetize cDNAs using oligo(dT) primers and affinity script reverse transcriptase (Agilent technologies France). Real-time quantitative PCR analyses were performed using the Light Cycler LC 1.5 (Roche, France). For each condition, expression was quantified in duplicate, and GAPDH or RNA18S was used as the housekeeping gene or normalizing RNA in the comparative cycle threshold (CT) method(Livak and Schmittgen, 2001) The sequences of primers used in this study are reported in supplemental data (Supplemental Data Table 1).

For the RNA sequencing, total RNA was isolated from 5 samples of LV from E16.5 mice and pooled either for Vitamin D depleted or control groups. The two pooled samples were used for the RNA-seq library preparation, using the kit TruSeq Stranded mRNA by Illumina.

Libraries were paired-end sequenced on the Illumina NextSeq 500 sequencer. Reads with a phred score lower than 20 and shorter than 25 bp were removed using Sickle (v1,33). Quality of trim reads was checked using multiQC (v1.0). Trim reads were aligned using STAR aligner (v2.7.0d) with arguments “outFilterMismatchNoverLmax” and “outFilterMultimapNmax” set to 0.08 and 1, respectively.

Transcripts discovery was performed using Cufflinks (v2.2.1) with the “library-type” argument set to fr-firstrand, and a GTF file obtained from GENCODE (“Comprehensive gene annotation”, vM1) provided as the genomic annotation. The GTF files produced for each sample by Cufflinks were combined using Cuffmerge. The “class code” assigned to each transcript by Cuffmerge was used to defined unknown transcripts (class code “u”). Only de novo transcripts with counts greater than 0 in at least one RNA-seq sample were kept for subsequent analyses. These de novo transcripts were combined with the GENCODE GTF file to produce the final genomic annotation that was provided to FeatureCounts (v1.6.1) for quantification.

Differential gene expression was performed using DESEQ2 between MVDD and CTRL. To create bigwig files, reads from Watson and Crick strands were selected using SAMtools (v1.9) and provided to the bam2wig.py script from the RseQC program suite (v2.6.4). RNA-seq profiles were visualized using the IGV genome browser. Data have been deposited to Genebank under http://www.ncbi.nlm.nih.gov/bioproject/682368

### Chromatin Immunoprecipitation (ChIP-sequencing)

Frozen embryonic left ventricles (from 8 hearts in each condition) at −80°C were thawed out and chromatin extracted. ChIP was performed using the anti-VDR antibody (Santacruz sc-13133) as previously described (Jebeniani et al., 2016).

Purified DNA was used for sequencing and QPCR. Primers used in QPCR are listed in supplemental table 2.

Fastq files were aligned to the mouse (mm10) reference genome, PCR duplicates were removed using Samtools and normalized genome coverage tracks were generated from uniquely mapping reads (mapq > 30) using deepTools2. Single-end reads, unmated reads, mate reads that map too far apart (> 4x fragment length) were extended by 200bp. Biological replicates were pooled and coverage was then calculated as average reads per million of mapped reads (RPM) in 10bp bins. To determine the peaks for ChIPs with narrow binding profiles, datasets were uniformly processed using the MACS2 with default parameters.

Data have been deposited to Genebank under https://www.ncbi.nlm.nih.gov/geo/query/acc.cgi?acc=GSE162895.

### ChIP-loop Chromosome Configuration Capture (3C)

Anti-CTCF ChIP-3-C was performed as previously described (Abboud et al., 2015; Jebeniani et al., 2016). Briefly, left ventricle of E16.5 embryonic hearts from control of VDD embryos were collected and freeze-clamped in liquid Nitrogen. The frozen hearts were smashed into powder in liquid Nitrogen, weighted, crosslinked with 1% formaldehyde for 10 min and quenched with 125 mM glycine for 5 min. The tissue was permeabilised and lysed as previously described (Jebeniani et al., 2016). Chromatin was digested overnight at 37°C by DpnII (New England Biolabs,. R0543M). The digestion was monitored by specific primers spanning DpnII sites in the genome (Supplemental Table 3)

CTCF was immunoprecipitated overnight at 4°C from 100 μg digested chromatin diluted in ChIP buffer (Jebeniani et al., 2016) using an anti-CTCF antibody (reference 07–729 Millipore France). The bound fraction was diluted and digested fragments ligated using T4-ligase (In vitrogen, 15224-041) for 4 hours at 16°C and then for 30 min at room temperature in the T4 ligase buffer. After a step of reverse-crosslink in the presence of proteinase K overnight at 65°C, and a 45 min incubation at 37°C with RNase (300 μg), DNA was extracted by phenol/ chloroform and 200 ng was used in real-time PCR. Amplicons were run on gel. DNA interaction frequencies at the CTCF sites were calculated as previously described (Abboud et al, 2015).

Primers used are listed in Supplemental table 3

### RNA-sequencing *Ingenuity Pathway Analysis (IPA)*

Differential gene expression (with p-value adjusted < 0.05), obtained from the RNA seq analysis between our conditions, were used in the IPA software to identify the canonical pathways differentially impacted by the maternal diet.

### Echocardiography

The cardiac function was evaluated using an Affiniti 50 ultrasound echocardiography (Philipps) equipped with a 11 MHz linear transducer (L12-4) on males at 2-months-old and 6-months old offspring. The animals were lightly anesthetized with inhaled isoflurane (1.5%) for maximum 30 min. Monitoring and analysis of echocardiography were carried out separately by at least two different persons.

### Statistical analysis

Data are expressed as mean ± SEM. Significant differences were determined by unpaired Student’s *t* test using GraphPad Prism. *P* < 0.05 was considered to be statistically significant.

---

**Supplemental table 1:**
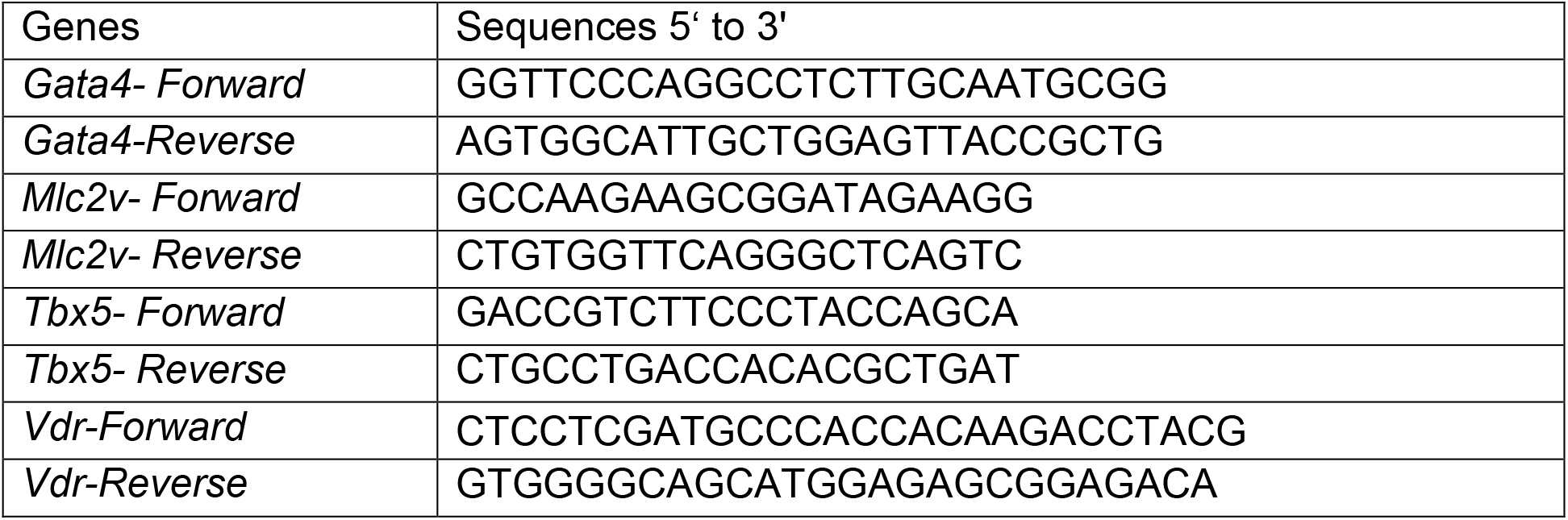
QPCR primers for genes

**Supplemental table 2:**
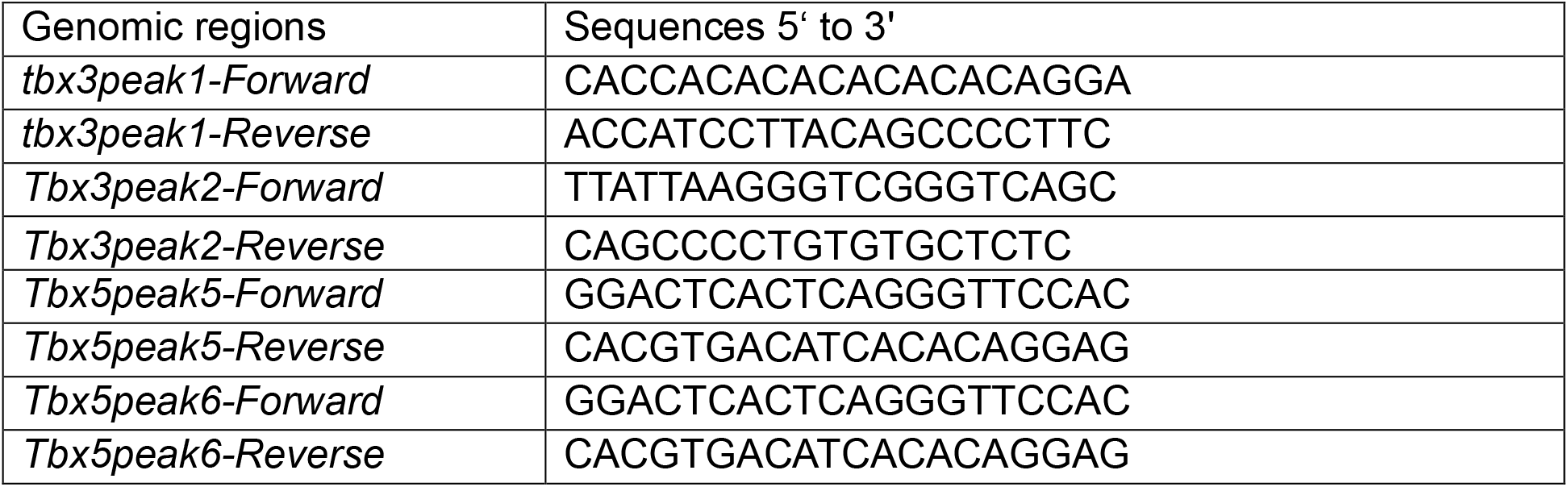
QPCR primers for anti-VDR ChIP-PCR

**Supplemental table 3:**
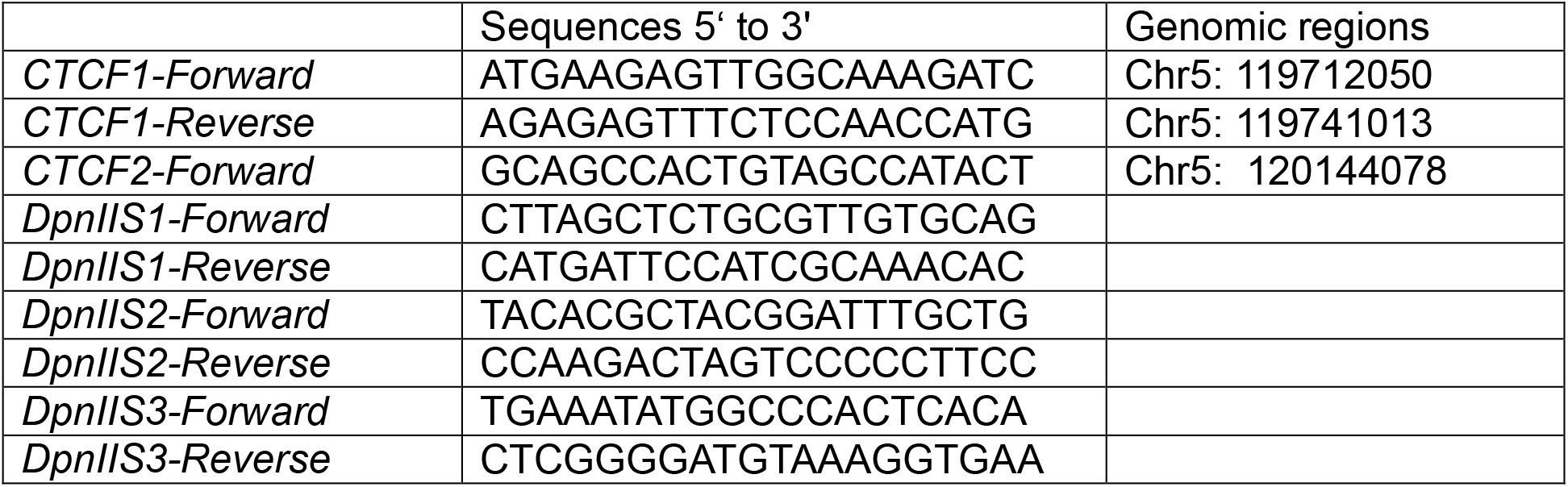
QPCR primers for anti-CTCF ChIP-loop

## References

Abboud, N., Moore-Morris, T., Hiriart, E., Yang, H., Bezerra, H., Gualazzi, M.G., Stefanovic, S., Guenantin, A.C., Evans, S.M., and Puceat, M. (2015). A cohesin-OCT4 complex mediates Sox enhancers to prime an early embryonic lineage. Nat Commun 6, 6749.

Assalin, H.B., Rafacho, B.P., dos Santos, P.P., Ardisson, L.P., Roscani, M.G., Chiuso-Minicucci, F., Barbisan, L.F., Fernandes, A.A., Azevedo, P.S., Minicucci, M.F., et al. (2013). Impact of the length of vitamin D deficiency on cardiac remodeling. Circ Heart Fail 6, 809–816.

Chen, T., Li, S.J., Chen, B., Huang, Q., Kong, X.Y., Shen, C., Gu, H.T., and Wang, X.W. (2019). Akt3 is a target of miR-29c-3p and serves an important function in the pathogenesis of congenital heart disease. Int J Mol Med 43, 980–992.

Christakos, S., Dhawan, P., Verstuyf, A., Verlinden, L., and Carmeliet, G. (2016). Vitamin D: Metabolism, Molecular Mechanism of Action, and Pleiotropic Effects. Physiol Rev 96, 365–408.

De Windt, L.J., Lim, H.W., Bueno, O.F., Liang, Q., Delling, U., Braz, J.C., Glascock, B.J., Kimball, T.F., del Monte, F., Hajjar, R.J., et al. (2001). Targeted inhibition of calcineurin attenuates cardiac hypertrophy in vivo. Proc Natl Acad Sci U S A 98, 3322–3327.

Fang, N., Hu, C., Sun, W., Xu, Y., Gu, Y., Wu, L., Peng, Q., Reiter, R.J., and Liu, L. (2020). Identification of a novel melatonin-binding nuclear receptor: Vitamin D receptor. J Pineal Res 68, e12618.

Gelb, B.D. (2015). History of Our Understanding of the Causes of Congenital Heart Disease. Circ Cardiovasc Genet 8, 529–536.

Georges, R., Nemer, G., Morin, M., Lefebvre, C., and Nemer, M. (2008). Distinct expression and function of alternatively spliced Tbx5 isoforms in cell growth and differentiation. Mol Cell Biol 28, 4052–4067.

Gezmish, O., Tare, M., Parkington, H.C., Morley, R., Porrello, E.R., Bubb, K.J., and Black, M.J. (2010). Maternal vitamin D deficiency leads to cardiac hypertrophy in rat offspring. Reprod Sci 17, 168–176.

Gong, L., Wang, S., Shen, L., Liu, C., Shenouda, M., Li, B., Liu, X., Shaw, J.A., Wineman, A.L., Yang, Y., et al. (2020). SLIT3 deficiency attenuates pressure overload-induced cardiac fibrosis and remodeling. JCI Insight 5.

Han, S., Li, T., Ellis, E., Strom, S., and Chiang, J.Y. (2010). A novel bile acid-activated vitamin D receptor signaling in human hepatocytes. Mol Endocrinol 24, 1151–1164.

Han, Y., Chen, A., Umansky, K.B., Oonk, K.A., Choi, W.Y., Dickson, A.L., Ou, J., Cigliola, V., Yifa, O., Cao, J., et al. (2019). Vitamin D Stimulates Cardiomyocyte Proliferation and Controls Organ Size and Regeneration in Zebrafish. Dev Cell 48, 853–863 e855.

Hanson, M.A., and Gluckman, P.D. (2014). Early developmental conditioning of later health and disease: physiology or pathophysiology? Physiol Rev 94, 1027–1076.

Hoffman, J.I., and Kaplan, S. (2002). The incidence of congenital heart disease. J Am Coll Cardiol 39, 1890–1900.

Jebeniani, I., Leschik, J., and Puceat, M. (2016). Epigenetic Regulation of Cardiac Differentiation of Embryonic Stem Cells and Tissues. J Vis Exp.

Kagiyama, S., Eguchi, S., Frank, G.D., Inagami, T., Zhang, Y.C., and Phillips, M.I. (2002). Angiotensin II-induced cardiac hypertrophy and hypertension are attenuated by epidermal growth factor receptor antisense. Circulation 106, 909–912.

Koster, M.P.H., van Duijn, L., Krul-Poel, Y.H.M., Laven, J.S., Helbing, W.A., Simsek, S., and Steegers-Theunissen, R.P.M. (2018). A compromised maternal vitamin D status is associated with congenital heart defects in offspring. Early Hum Dev 117, 50–56.

Kumar, S., Nanduri, R., Bhagyaraj, E., Kalra, R., Ahuja, N., Chacko, A.P., Tiwari, D., Sethi, K., Saini, A., Chandra, V., et al. (2020). Vitamin D3-VDR-PTPN6 axis mediated autophagy contributes to the inhibition of macrophage foam cell formation. Autophagy, 1–17.

Li, J., Kritzer, M.D., Michel, J.J., Le, A., Thakur, H., Gayanilo, M., Passariello, C.L., Negro, A., Danial, J.B., Oskouei, B., et al. (2013). Anchored p90 ribosomal S6 kinase 3 is required for cardiac myocyte hypertrophy. Circ Res 112, 128–139.

Livak, K.J., and Schmittgen, T.D. (2001). Analysis of relative gene expression data using real-time quantitative PCR and the 2(-Delta Delta C(T)) Method. Methods 25, 402–408.

Lu, M., Taylor, B.V., and Korner, H. (2018). Genomic Effects of the Vitamin D Receptor: Potentially the Link between Vitamin D, Immune Cells, and Multiple Sclerosis. Front Immunol 9, 477.

Luppino, J.M., Park, D.S., Nguyen, S.C., Lan, Y., Xu, Z., Yunker, R., and Joyce, E.F. (2020). Cohesin promotes stochastic domain intermingling to ensure proper regulation of boundary-proximal genes. Nat Genet 52, 840–848.

Mokhtar, W.A., Fawzy, A., Allam, R.M., Amer, R.M., and Hamed, M.S. (2019). Maternal vitamin D level and vitamin D receptor gene polymorphism as a risk factor for congenital heart diseases in offspring; An Egyptian case-control study. Genes Dis 6, 193–200.

Moore-Morris, T., van Vliet, P.P., Andelfinger, G., and Puceat, M. (2018). Role of Epigenetics in Cardiac Development and Congenital Diseases. Physiol Rev 98, 2453–2475.

Nagpal, S., Na, S., and Rathnachalam, R. (2005). Noncalcemic actions of vitamin D receptor ligands. Endocr Rev 26, 662–687.

Parelho, V., Hadjur, S., Spivakov, M., Leleu, M., Sauer, S., Gregson, H.C., Jarmuz, A., Canzonetta, C., Webster, Z., Nesterova, T., et al. (2008). Cohesins functionally associate with CTCF on mammalian chromosome arms. Cell 132, 422–433.

Paschou, S.A., Kosmopoulos, M., Nikas, I.P., Spartalis, M., Kassi, E., Goulis, D.G., Lambrinoudaki, I., and Siasos, G. (2019). The Impact of Obesity on the Association between Vitamin D Deficiency and Cardiovascular Disease. Nutrients 11.

Rahman, A., Hershey, S., Ahmed, S., Nibbelink, K., and Simpson, R.U. (2007). Heart extracellular matrix gene expression profile in the vitamin D receptor knockout mice. J Steroid Biochem Mol Biol 103, 416–419.

Reddy Vanga, S., Good, M., Howard, P.A., and Vacek, J.L. (2010). Role of vitamin D in cardiovascular health. Am J Cardiol 106, 798–805.

Sales, V.M., Ferguson-Smith, A.C., and Patti, M.E. (2017). Epigenetic Mechanisms of Transmission of Metabolic Disease across Generations. Cell Metab 25, 559–571.

Saraf, R., Morton, S.M., Camargo, C.A., Jr., and Grant, C.C. (2016). Global summary of maternal and newborn vitamin D status - a systematic review. Matern Child Nutr 12, 647–668.

Seipelt, E.M., Tourniaire, F., Couturier, C., Astier, J., Loriod, B., Vachon, H., Puceat, M., Mounien, L., and Landrier, J.F. (2020). Prenatal maternal vitamin D deficiency sex-dependently programs adipose tissue metabolism and energy homeostasis in offspring. FASEB J.

Seuter, S., Neme, A., and Carlberg, C. (2016). Epigenome-wide effects of vitamin D and their impact on the transcriptome of human monocytes involve CTCF. Nucleic Acids Res 44, 4090–4104.

Takeuchi, J.K., Ohgi, M., Koshiba-Takeuchi, K., Shiratori, H., Sakaki, I., Ogura, K., Saijoh, Y., and Ogura, T. (2003). Tbx5 specifies the left/right ventricles and ventricular septum position during cardiogenesis. Development 130, 5953–5964.

van Weerd, J.H., Badi, I., van den Boogaard, M., Stefanovic, S., van de Werken, H.J., Gomez-Velazquez, M., Badia-Careaga, C., Manzanares, M., de Laat, W., Barnett, P., et al. (2014). A large permissive regulatory domain exclusively controls Tbx3 expression in the cardiac conduction system. Circ Res 115, 432–441.

Wang, T.J., Pencina, M.J., Booth, S.L., Jacques, P.F., Ingelsson, E., Lanier, K., Benjamin, E.J., D’Agostino, R.B., Wolf, M., and Vasan, R.S. (2008). Vitamin D deficiency and risk of cardiovascular disease. Circulation 117, 503–511.

Xiang, W., Kong, J., Chen, S., Cao, L.P., Qiao, G., Zheng, W., Liu, W., Li, X., Gardner, D.G., and Li, Y.C. (2005). Cardiac hypertrophy in vitamin D receptor knockout mice: role of the systemic and cardiac renin-angiotensin systems. Am J Physiol Endocrinol Metab 288, E125–132.

Zaidi, S., and Brueckner, M. (2017). Genetics and Genomics of Congenital Heart Disease. Circ Res 120, 923–940.

