## supplemental figures for "Maternal vitamin D deficiency impairs heart formation in mouse offspring through a change in 3D-chromatin structure"

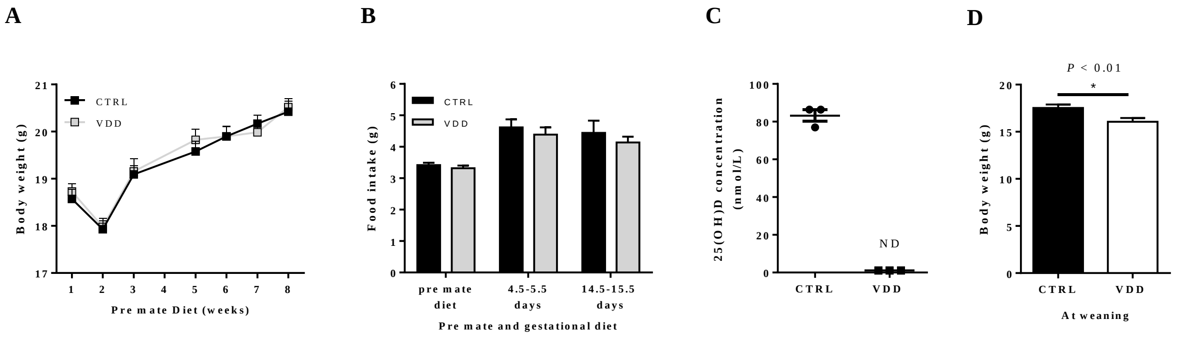


Supplemental Data Figure 1. General parameters of maternal model, and juvenile offspring. (A) Growing curve of the female during the 8-weeks of pre-mate diet (i.e. CTRL or VDD). (B) Maternal food intake during pre-mate and gestational diets. (C) Maternal 25-dihydroxyvitamin D plasma concentration. (D) Body weight of male at weaning regarding maternal diet ($P < 0.01$)

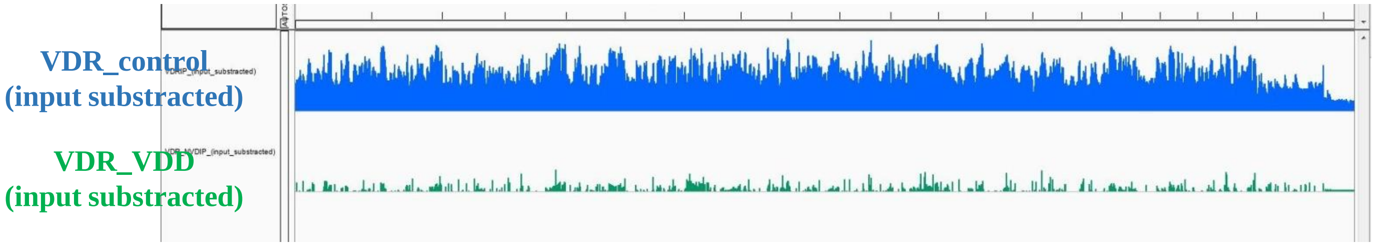


Supplemental Data Figure 2. whole genome anti-VDR ChIP-sequencing

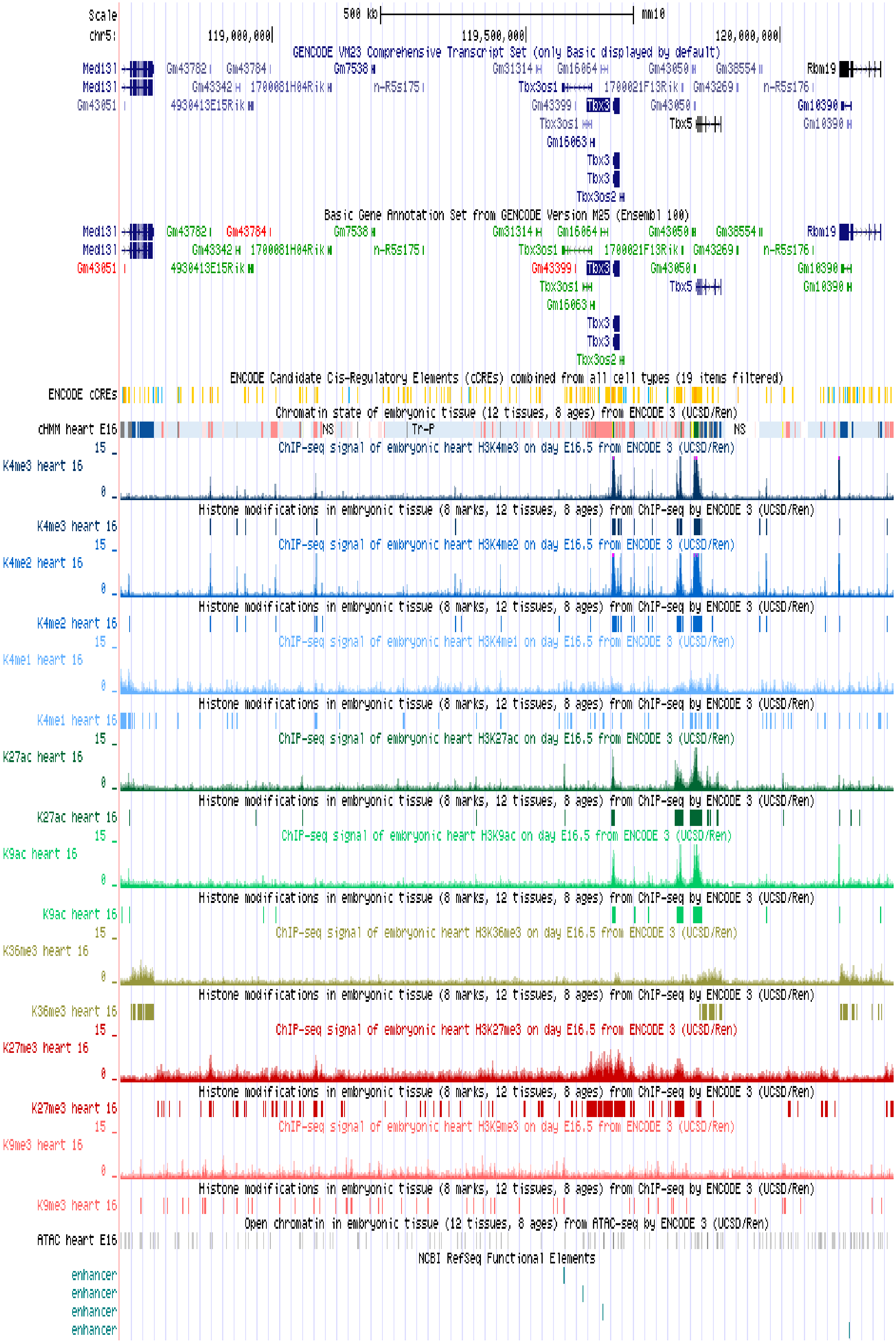


Supplemental Data Figure 3: Encode data Epigenetic marks within the *Tbx3/Tbx5* loci
